# Tamoxifen Suppresses Pancreatic β-Cell Proliferation in Mice

**DOI:** 10.1101/586602

**Authors:** Surl-Hee Ahn, Anne Granger, Matthew M. Rankin, Carol J. Lam, Aaron R. Cox, Jake A. Kushner

## Abstract

Tamoxifen is a mixed agonist/antagonist estrogen analogue that is frequently used to induce conditional gene deletion in mice using Cre-loxP mediated gene recombination. Tamoxifen is routinely employed in extremely high-doses relative to typical human doses to induce efficient gene deletion in mice. Although tamoxifen has been widely assumed to have no influence upon β-cells, the acute developmental and functional consequences of high-dose tamoxifen upon glucose homeostasis and adult β-cells are largely unknown. We tested if tamoxifen influences glucose homeostasis in male mice of various genetic backgrounds. We then carried out detailed histomorphometry studies of mouse pancreata. We also performed gene expression studies with islets of tamoxifen-treated mice and controls. Tamoxifen had modest effects upon glucose homeostasis of mixed genetic background (F1 B6129SF1/J) mice, with fasting hyperglycemia and improved glucose tolerance but without overt effects on fed glucose levels or insulin sensitivity. Tamoxifen inhibited proliferation of β-cells in a dose-dependent manner, with dramatic reductions in β-cell turnover at the highest dose (decreased by 66%). In sharp contrast, tamoxifen did not reduce proliferation of pancreatic acinar cells. β-cell proliferation was unchanged by tamoxifen in 129S2 mice but was reduced in C57Bl6 genetic background mice (decreased by 59%). Gene expression studies revealed suppression of RNA for cyclins D1 and D2 within islets of tamoxifen-treated mice. Tamoxifen has a cytostatic effect on β-cells, independent of changes in glucose homeostasis, in mixed genetic background and also in C57Bl6 mice. Tamoxifen should be used judiciously to inducibly inactivate genes in studies of glucose homeostasis.

## INTRODUCTION

β-cell regeneration is a fundamental goal in diabetes research that will require elucidation of the numerous circulating and intrinsic signals that govern growth and expansion of mature adult β-cells, such as glucose, hormones, and various growth factors [1]. Adult β-cells are largely the products of self-renewal [2,3]. Self-renewal capacity of adult β-cells is limited by a replication refractory period that prevents cells that have recently divided from immediately reentering the cell cycle [3-5]. In addition, β-cell regenerative capacity decreases with age by largely unknown mechanisms [1,6-8]. The aging process has been commonly linked with cellular senescence, a state in which a cell no longer has the ability to proliferate, and limited regenerative capacity. In response to increased metabolic requirements such as obesity, pregnancy, and after pancreatic injury like partial pancreatectomy, shortening of the refractory period allows for a compensatory increase in β-cell mass [3,5,9]. Failure to adapt to increased metabolic demand results in insulin deficiency and type 1 and type 2 diabetes. Understanding the mechanisms that govern β-cell proliferation could be crucial in restoring β-cell mass in patients with diabetes.

Inducible gene deletion has emerged as a widely employed tool to interrogate developmental signals in mice. Early studies employed germline gene deletions, revealing that many genes had primordial roles that were essential for early embryonic development. As a result, it was difficult to ascertain the roles of these genes in postnatal life [10]. Many genes were active in multiple tissues of interest, which further complicated studies to determine the tissue-specific roles of specific gene products in particular tissues of interest. Consequently, tissue-specific inducible approaches arose as a popular approach to determine the roles of genes.

Tissue-specific inducible gene deletion is often used to determine the function of various genes or cells of origin in the formation, maintenance, or regeneration of β-cells. Although tetracycline inducible methods have been occasionally employed, a more common approach for β-cell-specific gene deletion involves the anticancer drug tamoxifen and transgenic mice that express a tamoxifen-dependent form of Cre recombinase in β-cells or their progenitors. In this system Cre is fused to a mutated ligand-binding domain of the human estrogen receptor (ER) [11]. This tool allows temporal control and tissue-specific gene deletion during embryonic or postnatal development, or in adult life. The tamoxfen-mutant estrogen receptor ligand-binding domain (ERT) preferentially binds to tamoxifen and is insensitive to endogenous estrogen [12]. Fusion of ERT to Cre allows tamoxifen-inducible Cre (Cre^ERT^) which is free to enter the nucleus and therefore induce a targeted mutation at loxP sites [13]. The Cre^ERT^-loxP can inducibly and specifically target loxP sites within tissues that express a particular transgenic promoter. Generation of inducible Cre lines driven by *Ins1, Ins2*, and *Pdx1* promoters has allowed numerous groups to study adult β-cell biology, bypassing embryonic lethal gene inactivation or more precisely discerning tissue-specific effects [14,15].

Estrogens and tamoxifen have profound effects on physiology throughout the body. Estrogen receptors have functionally independent DNA- and ligand-binding domains that activate or repress transcription upon binding of the ligand [16]. Estrogen receptors (ERα and ERβ) are also highly sophisticated molecular switches that are capable of responding to two cellular signaling pathways, simultaneously transducing ligand bound and also membrane-associated receptors signals [16]. Estrogens have widespread systemic effects on glucose homeostasis, β-cells, and obesity [17,18]. As an estrogen receptor mixed agonist/antagonist, tamoxifen may also influence cell cycle entry and has been shown to have potent effects on proliferation and/or apoptosis of tumor cells via estrogen receptors ERα and ERβ [19]. In addition, at high concentrations tamoxifen-mediated growth inhibition may act independent of ERs by inhibition of protein kinase C (PKC) activity [20,21].

Despite widespread use of tamoxifen to induce gene recombination, the dose-response relationship of tamoxifen on β-cell turnover has not been studied in non-diabetic mice. Given the published cytostatic and cytotoxic effect of tamoxifen on tumor cells, we sought to investigate potential effects of tamoxifen on β-cell turnover. Here, we show that tamoxifen directly inhibits β-cell proliferation in adult mice, without affecting cell survival and independent of any changes in glucose homeostasis.

## RESEARCH DESIGN AND METHODS

### Mice

All experiments with mice were performed in the animal facility at Children’s Hospital of Philadelphia, under the supervision of and according to the guidelines of the Institutional Animal Care and Use Committee. The IACUC prospectively approved this research. CHOP IACUC #656. Male F1 hybrid B6129SF1/J mice (stock 101043) were obtained at 2 months of age from Jackson Laboratory (Bar Harbor, ME). Inbred C57Bl/6J (Jackson stock 000664) and 129S2 mice (stock 129S2/SvPasCrl) were purchased from Jackson Laboratory and Charles River Laboratories (Wilmington, MA), respectively. Mice labeled continuously with 5-bromo-2-deoxyuridine (BrdU) in the drinking water, as previously [22]. Euthanasia was performed via pentobarbital overdose (60 mg/kg).

### Tamoxifen

Tamoxifen citrate salt from Sigma-Aldrich (St. Louis, Mo) was dissolved in 0.2ml 10% Etoh in corn oil and administered via oral gavage for five days. Control animals received oral gavage of 0.2ml 10%EtoH in oil.

### Glucose homeostasis

Intraperitoneal glucose tolerance tests were performed on mice fasted for 16 hours with 2 g D-glucose per kg body weight as previously [23]. Insulin sensitivity tests were performed on mice fasted for 4 hours. The mice were injected with 0.75 U per kg body weight human regular insulin (diluted to 0.075 U/mL). Blood glucose was measured at 0,15, 30, 60, and 120 minutes after insulin injection.

### Pancreatic Morphometry

β-cell morphometry was performed with Volocity 6.1.1 (PerkinElmer, Waltham, MA)[24,25]. A Zeiss AxioImager (Carl Zeiss, Thornwood, NY) with automated X-Y stage and Orca ER camera (Hamamatsu, Middlesex, NJ) acquired thousands of islet images with tens of thousands of individual nuclei analyzed/sample, as previously [23]. To quantify β-cell area and mass, all possible β-cells were quantified from 3-4 lateral pancreatic sections for both head and tail. Primary antisera: guinea pig anti-insulin (Zymed/Invitrogen).

### Cell proliferation

Mice were continuously labeled with 5-bromo-2′-deoxyuridine (BrdU) for 14 days after gavage before they were euthanized. Paraformaldehyde-fixed, paraffin-embedded sections were stained with DAPI/insulin/BrdU or DAPI/insulin/BrdU/Ki67. Images were acquired from 20-30 islets per animal and condition, which represented 2,500–4,000 β-cells per animal, as previously [23,26,27].

### Cell death

Apoptosis analysis was performed as previously [28]. ∼ 10,000 β-cells per condition were analyzed for the total intra-islet TUNEL (terminal deoxynucleotidyl-transferase-mediated dUTP nick end labeling) positive cells.

### Islet Isolation and real-time quantitative PCR (RT-PCR)

Islets were isolated from mice 14 days after tamoxifen treatment (with 0.3mg/g body weight) and processed into cDNA, as previously [7,29,30]. Groups of 30 to 40 islets were used for gene expression studies. Real-time quantitative dual fluorescent–labeled fluorescence resonance energy transfer (FRET) PCR (50°C for 2 min and 95°C for 10 min, followed by 40 cycles of 95°C for 15 s and 60°C for 1 min) was performed to amplify triplicate samples, with primer-probe sets as previously [7,29,30]. Relative gene product amounts were reported for each gene compared with cyclophilin. Results are reported from a single experiment with biological repeats averaged and reported as means ± SEM.

### Statistics

All results are reported as means ± SEM for equivalent groups. Results were compared with independent Student’s t-tests (unpaired and two-tailed) reported as P values.

## RESULTS

### Tamoxifen is well-tolerated in mixed genetic background mice

To assess the potential effects of tamoxifen in vivo, we carried out a treatment study with two-month-old hybrid male mice (F1 hybrid B6129SF1/J). We used this strain because they closely resemble the mixed genetic background typical of genetic knockout mice. We choose to focus on male mice to minimize the impact of estrus cycle on our studies. One group was designated as a control and gavaged with vehicle (oil), while the tamoxifen-treated groups were gavaged with drug (0.025, 0.1, 0.2, or 0.3mg/g body weight) for 5 days (Fig. 1a). The range of tamoxifen doses was chosen to approach a maximum of 8mg per day, the typical amount employed to induce gene deletion when studying glucose homeostasis or β-cell lineage relationships in adult mice [2]. Consistent with previous reports, tamoxifen-treated mice were viable and phenotypically healthy during the course of studies. Body weight of the mice was not altered by tamoxifen treatment at any dose (Fig. 1b, Table S1).

**Fig. 1.**
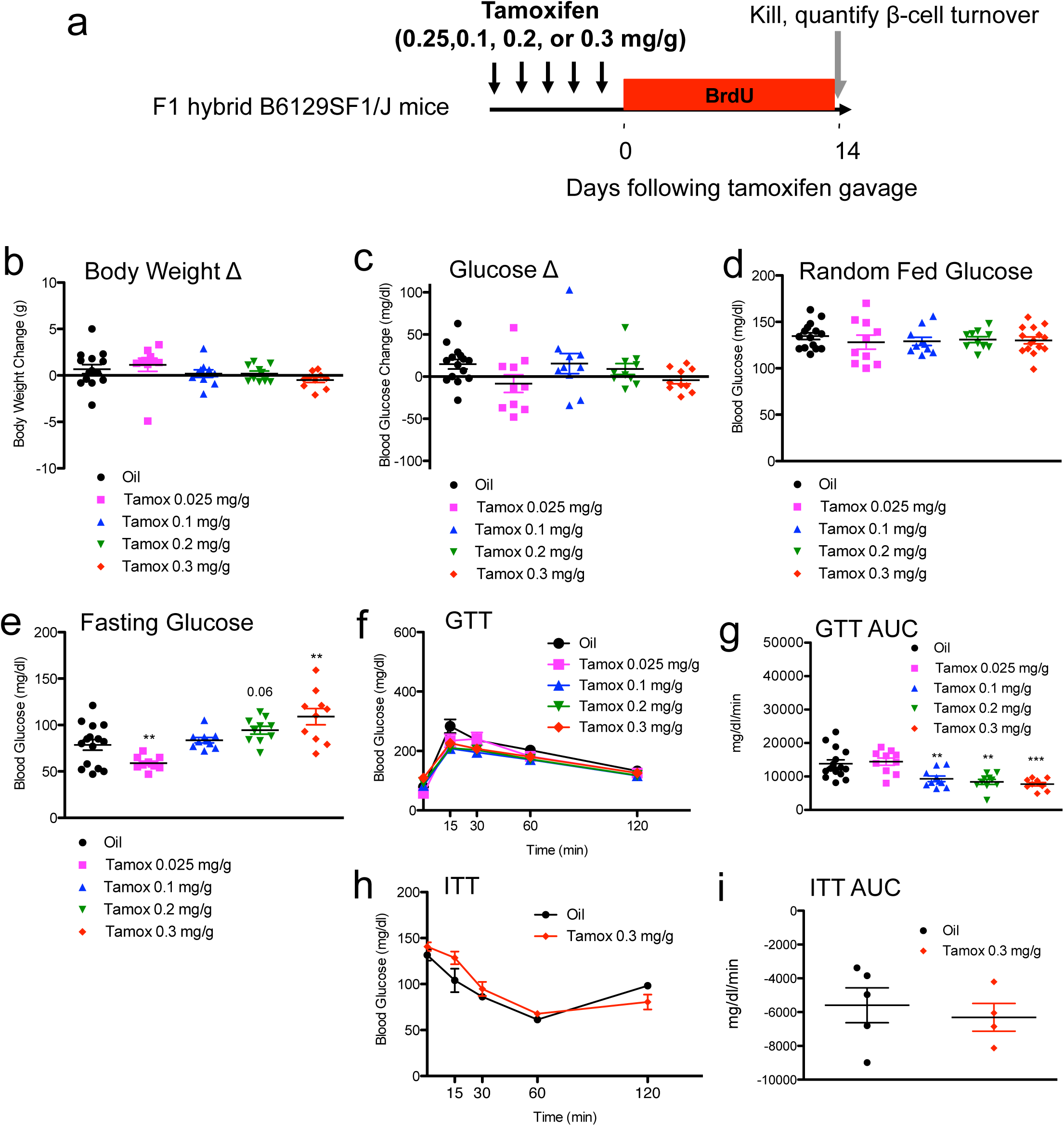
Tamoxifen increases fasting glucose but improves glucose tolerance in mixed genetic background (F1 hybrid B6129SF1/J) mice. (**a**) Schematic indicating time course of treatment. Mice received daily oral gavage with tamoxifen (0.025, 0.1, 0.2, or 0.3mg/g body weight) or vehicle for 5 days. Mice were sequentially labeled with BrdU for two weeks each, starting immediately after tamoxifen treatment. (**b, c**) Body weight (**b**) and blood glucose (**c**) change from the start of treatment (day 0) to the end of the experiment (day 19). Random fed (**d**) and fasting (**e**) blood glucose levels at the end of the experiment. (**f, g**) Glucose tolerance tests (GTT) with glucose curves (**f**) and GTT area under curve (AUC) calculations (**g**). (**h, i**) Insulin tolerance tests (GTT) with glucose curves (**h**) and ITT area under curve (AUC) calculations (**i**). Results expressed as mean ± SEM for 10 mice per group. **p≤0.01, ***p≤0.001, tamoxifen vs. vehicle.

### Tamoxifen influences glucose homeostasis in mixed genetic background mice

We tested for alteration of glucose homeostasis in the tamoxifen-treated mice. Reassuringly, random fed blood glucose did not change with tamoxifen treatment (Fig. 1c, Table S1). Similarly, random fed glucose was not significantly different within tamoxifen-treated groups at the end of the study (Fig. 1d, Table S1). Interestingly, fasting blood glucose was moderately decreased in the low-dose (0.025 mg/d) cohort compared to controls (Fig. 1e, Table S1). In contrast, high-dose (0.2 or 0.3 mg/d) mice actually displayed increased fasting blood glucose compared to controls (Fig. 1e, Table S1).

We performed provocative testing of glucose homeostasis in the tamoxifen-treated mixed genetic background mice. Glucose tolerance was equivalent in the low dose (0.025 mg/d) cohort (Fig. 1f, Table S1). However, glucose tolerance was significantly improved in each of the high-dose tamoxifen-treated cohorts when compared to controls. As tamoxifen has been shown to influence peripheral insulin action [31-33] we also performed insulin sensitivity tests. But, high-dose tamoxifen (0.3 mg/g body weight) had no discernable effect upon insulin sensitivity compared to controls (Fig. 1e, Table S1). In summary, high-dose tamoxifen was associated with increased fasting glucose but also appeared to have moderately beneficial effects upon glucose tolerance.

### β-cell morphology

We characterized pancreatic histology in the tamoxifen-treated mice. As expected, β-cell morphology was fully intact, without obvious deficits upon β-cells or islet endocrine cells. Pancreatic exocrine histology was similarly normal in appearance, without any evidence of pancreatitis, acinar destruction, lymphocytic infiltrates, or ductal hyperplasia. We used high throughput imaging methodologies to measure β-cell content, quantifying total pancreas and insulin area in high-dose tamoxifen-treated mice (0.3 mg/g) and controls. Pancreas mass was slightly reduced in tamoxifen-treated mice compared to controls (Table S2). Tamoxifen-treated mice also exhibited non-significant reductions in β-cell area (reduced by 17%, p=0.43) and β-cell mass (reduced by 29%, p=0.20)(Figure 2a-b, Table S2).

**Fig. 2.**
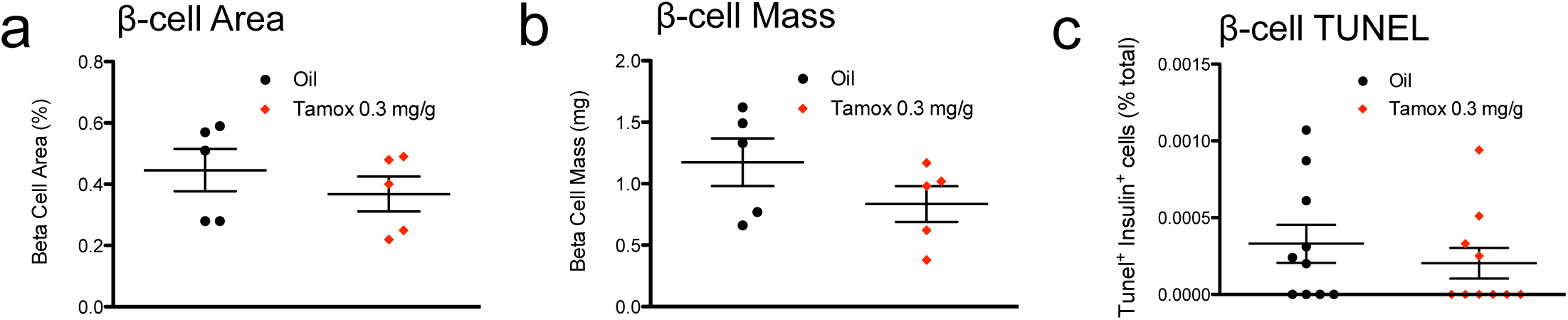
Tamoxifen does not influence β-cell area, mass or survival in mixed genetic background mice. (**a-b**) β-cell morphometry, as quantified by β-cell area (**a**) or β-cell mass (**b**). Results expressed as mean ± SEM for 5 mice per group. (**c**) β-cell survival, as quantified by intra-islet TUNEL cells over total islet β-cells. Results expressed as mean ± SEM for 10 mice per group.

### Tamoxifen does not alter β-cell survival

Estrogen-related signals have been widely reported to influence β-cell survival in pathological conditions [34-41]. As a result, we considered the possibility that acute effects of high-dose tamoxifen administration via ERα blockade (or some other target) might adversely impact β-cell survival. We quantified β-cell death via TUNEL labeling in pancreata from tamoxifen-treated mice and controls. Reassuringly, pancreas sections from high-dose tamoxifen mice and controls contained characteristically low levels of apoptotic β-cells (Fig. 2c, Table S3). We cannot rule out the possibility that tamoxifen might have subtly impaired β-cell survival in our cohort. Nevertheless, our TUNEL studies are consistent with earlier studies illustrating the lack of effect on β-cell mass from tamoxen, Thus, high-dose tamoxifen did not seem to grossly impair β-cell survival in healthy non-diabetic mice.

### Tamoxifen inhibits β-cell proliferation in a dose-dependent and tissue-specific manner

Mice were continuously labeled with BrdU for 2 weeks in the drinking water to capture all β-cell replicative events and then euthanized. Islets exhibited normal morphology in all groups, unaltered by any dose of tamoxifen (Fig. 3a-e). To test if tamoxifen altered β-cell proliferation we quantified BrdU and insulin co-positive cells in the pancreata. β-cell proliferation was progressively inhibited by increasing doses of tamoxifen: β-cell BrdU was decreased by 12% in mice treated with 0.025 mg/g (p=0.19, n=10), by 24% with 0.1 mg/g (p=0.049, n=10), by 41% with 0.2 mg/g (p=0.0002, n=10), and by 66% with 0.3mg/g tamoxifen (p=0.00000006, n=10)(Fig. 3f, Table S4). These studies reveal that tamoxifen reduced β-cell proliferation in a dose-dependent manner in mixed genetic background mice.

**Fig. 3.**
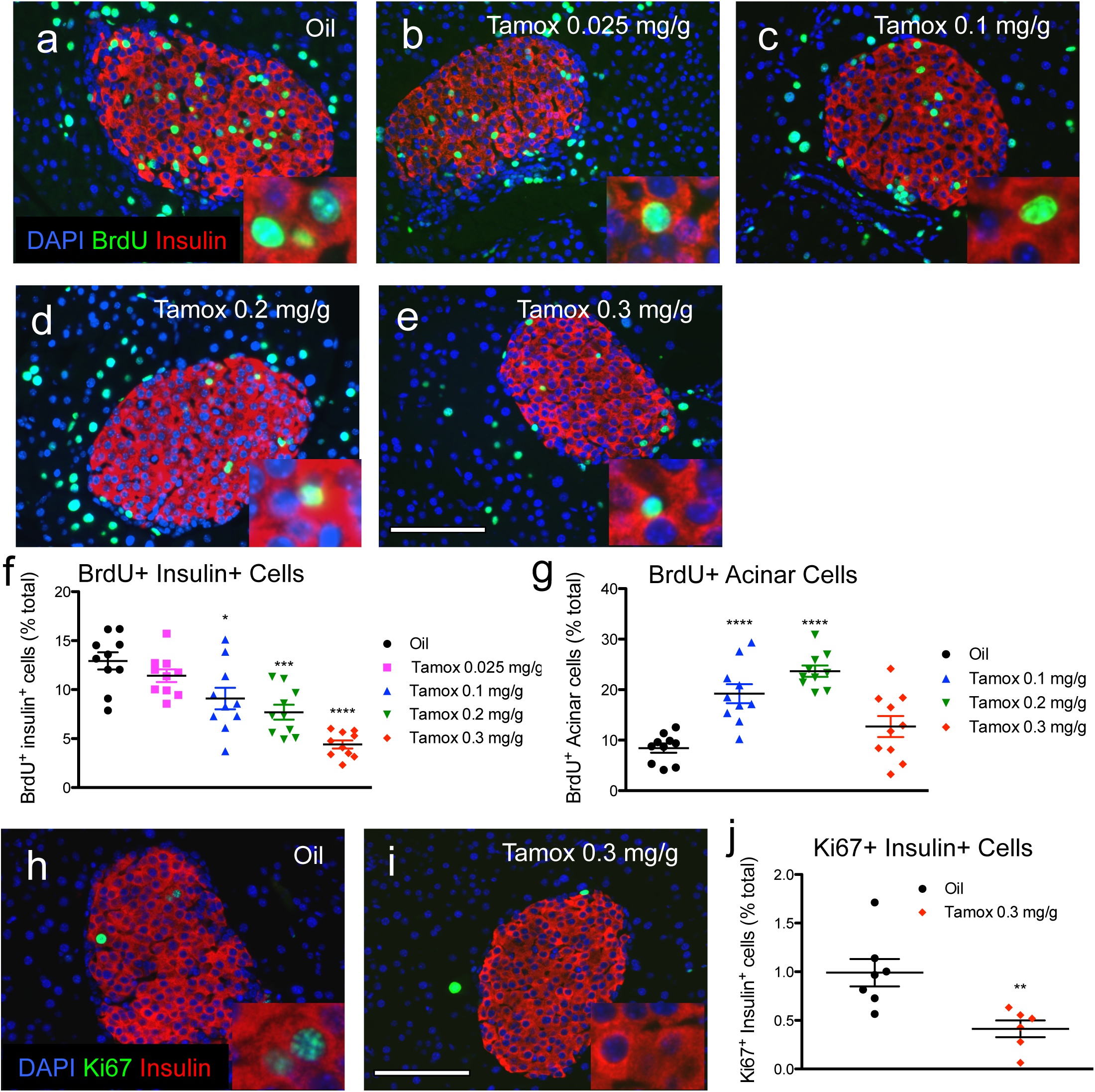
Tamoxifen reduces β-cell proliferation in mixed genetic background mice. (**a-e**) Representative islet images for vehicle (**a**) and increasing doses of tamoxifen (**b-e**) stained for insulin (red), BrdU (green), and DAPI (blue). Scale bar: 100µm. (**c**) Cumulative β-cell proliferation of BrdU+ insulin+ cells after continuous labeling with BrdU 2 weeks following tamoxifen treatment. (**d**) Cumulative acinar cell proliferation of BrdU+ acinar cells. Results expressed as mean ± SEM for 10 mice per group. (**h, i**) Representative islet images for vehicle (**h**) and tamoxifen (**i**) stained for insulin (red), Ki67 (green), and DAPI (blue). Scale bar: 100µm. (**j**) β-cell proliferation of Ki67+ insulin+ cells. *p≤0.05, **p≤0.01, ***p≤0.001, ****p≤0.001, tamoxifen vs. vehicle.

To test if tamoxifen had generalized effects upon pancreatic cell division we quantified BrdU incorporation into acinar tissue. In contrast to our results with β-cells, acinar cell proliferation was actually increased in the tamoxifen-treated cohorts: by 130% with 0.1 mg/g (p= 0.00007), by 180% with 0.2 mg/g (p= 0.000000003), and by 50% with 0.3mg/g tamoxifen (p= 0.13)(Fig. 3g, Table S4). Thus, tamoxifen seemed to increase pancreatic acinar cell proliferation, in sharp contrast to tamoxifen inhibition of β-cell proliferation.

We performed Ki67 immunostaining to further quantify β-cell proliferation in pancreata from tamoxifen-treated mice. Ki67 is specific to dividing cells and is therefore useful to detect active β-cell proliferation. In contrast, continuous BrdU labeling integrates β-cell proliferation over the entire labeling period (2 weeks). Detection of Ki67 in β-cells at the time of tissue harvest (2 weeks after the end of tamoxifen treatment) allowed us to test whether the tamoxifen-mediated growth inhibition of β-cells was only temporary. Interestingly, β-cell proliferation, measured by insulin+ Ki67+ cells, was still reduced (by 59%) in animals treated with 0.3 mg/g tamoxifen at the end of the experiment (p=0.04, n=5-7) (Fig. 3h-j, Table S5). These results confirm our BrdU incorporation findings of reduced β-cell proliferation in tamoxifen-treated mice. In addition, the Ki67 results also indicate that tamoxifen has a lasting impact, with depressed β-cell proliferation notable even after a 2 week washout recovery period.

### Tamoxifen inhibition of β-cell proliferation is genetic background-dependent

We carried out studies to test to determine genetic background-specific effects of tamoxifen upon glucose homeostasis and β-cell proliferation. We treated a cohort of pure 129S2 mice with tamoxifen (0.2 mg/g body weight) or oil and tested for defects in glucose homeostasis (Fig. 4a). We choose this intermediate dose of tamoxifen to minimize the chance of severe side effects in this pure strain that might threaten mouse viability. Similar to our mixed genetic background studies, 129S2 mice tolerated tamoxifen well, with no effect upon body weight or random fed glucose levels (Fig. 4b-d, Table S6). Similar to mixed genetic background mice, fasting glucose was moderately increased compared to controls (Fig. 4e, Table S6). Glucose tolerance was slightly improved 5 days post tamoxifen but not at 14 days (Fig. 4f-i, Table S6). The glucose homeostasis phenotypes of tamoxifen-treated 129S2 mice resemble that of tamoxifen-treated F1 hybrid B6129SF1/J mice, with moderate fasting hyperglycemia (Fig. 1e) and improved glucose tolerance (Fig. 1g).

**Fig. 4.**
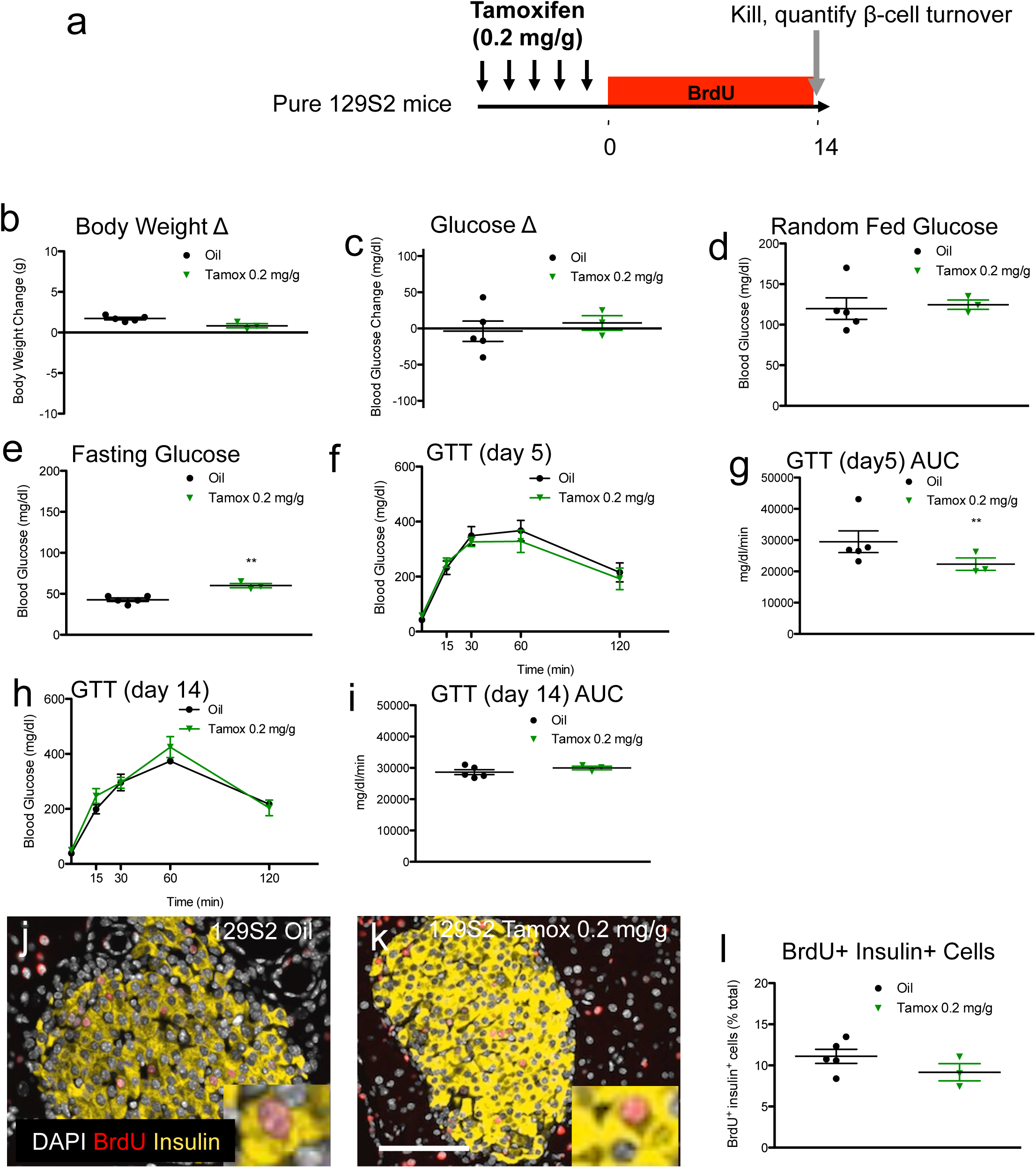
Tamoxifen increases fasting glucose but improves glucose tolerance in pure 129SF1/J mice; Tamoxifen does not significantly reduce β-cell proliferation in pure 129SF1/J mice. (**a**) Schematic indicating time course of treatment. 129SF1/J mice received daily oral gavage with tamoxifen (0.2mg/g body weight) or vehicle for 5 days. Mice were sequentially labeled with BrdU for two weeks each, starting immediately after tamoxifen treatment. (**b, c**) Body weight (**b**) and blood glucose (**c**) change from the start of treatment (day 0) to the end of the experiment (day 19). Random fed (**d**) and fasting (**e**) blood glucose levels at the end of the experiment. (**f,g**) GTT curves (**f**) and GTT AUC calculations (**g**) for 5 days after tamoxifen treatment. (**h, i**) GTT curves (**h**) and GTT AUC (**i**) calculations for 5 days after tamoxifen treatment. (**j, k**) Representative islet images for vehicle (**j**) and tamoxifen (**k**) stained for insulin (red), BrdU (green), and DAPI (blue). Scale bar: 100µm. (**l**) Cumulative β-cell proliferation of BrdU+ insulin+ cells after continuous labeling with BrdU 2 weeks following tamoxifen treatment. Results expressed as mean ± SEM for 5 controls and 3 tamoxifen-treated mice per group. **p≤0.01, tamoxifen vs. vehicle.

We tested for genetic background-specific effects of tamoxifen on β-cell proliferation in the 129S2 mice. As previously, mice were continuously labeled with BrdU for 2 weeks in the drinking water (Fig. 4a). As expected, tamoxifen-treated 129S2 mice exhibited normal islet morphology (Fig. 4j-k). In contrast to the mixed genetic background mice, tamoxifen-treated 129S2 mice only exhibited slight (∼17%) and non-significant (p=0.21) trends to reduced β-cell proliferation compared to controls (Fig. 4l, Table S7).

We also treated a cohort of pure C57Bl/6J mice with tamoxifen (Fig. 5a). C57Bl/6J mice also tolerated tamoxifen well, without any discernable impact on weight or random glucose levels (Fig. 5b-d, Table S6). In contrast to the hybrid or 129S2 mice, fasting glucose in the C57Bl/6J mice was actually reduced compared to controls (Fig. 5e, Table S6). At 5 days and 14 days post tamoxifen glucose tolerance was only slightly (11 and 17%) and non-significantly (p=0.22 and 0.22) improved (Fig. 5f-i, Table S6).

**Fig. 5.**
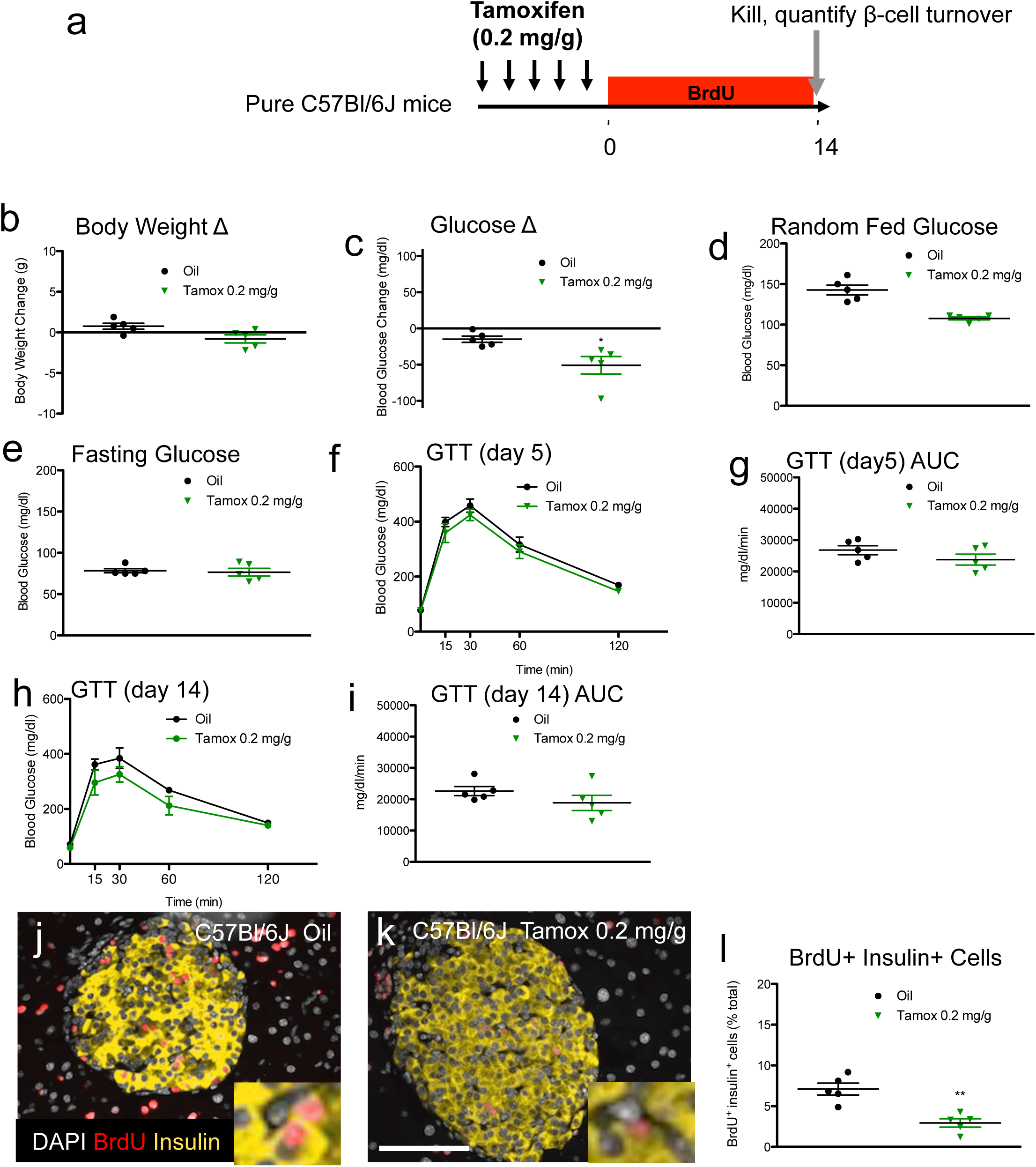
Tamoxifen increases fasting glucose but improves glucose tolerance in C57Bl/6J mice; Tamoxifen significantly reduces β-cell proliferation in C57Bl/6J mice. (**a**) Schematic indicating time course of treatment. C57Bl/6J mice received daily oral gavage with tamoxifen (0.2mg/g body weight) or vehicle for 5 days. Mice were sequentially labeled with BrdU for two weeks each, starting immediately after tamoxifen treatment. (**b, c**) Body weight (**b**) and blood glucose (**c**) change from the start of treatment (day 0) to the end of the experiment (day 19). Random fed (**d**) and fasting (**e**) blood glucose levels at the end of the experiment. (**f, g**) GTT curves (**f**) and GTT AUC calculations (**g**) for 5 days after tamoxifen treatment. (**h, i**) GTT curves (**h**) and GTT AUC (**i**) calculations for 5 days after tamoxifen treatment. (**j, k**) Representative islet images for vehicle (**j**) and tamoxifen (**k**) stained for insulin (red), BrdU (green), and DAPI (blue). Scale bar: 100µm. (**l**) Cumulative β-cell proliferation of BrdU+ insulin+ cells after continuous labeling with BrdU 2 weeks following tamoxifen treatment. Results expressed as mean ± SEM for 5 controls and 5 tamoxifen-treated mice per group. **p≤0.01, tamoxifen vs. vehicle.

As expected, tamoxifen-treated C57Bl/6J mice exhibited normal islet morphology (Fig. 5j-k). But, β-cell proliferation was dramatically reduced in the tamoxifen cohort compared to controls (by 59%, p=0.002) (Fig. 5l, Table S7). Thus, tamoxifen selectively inhibits β-cell proliferation in C57Bl/6J mice but not in 129S2 mice. Notably, the reduced β-cell proliferation phenotype of the C57Bl/6J mice occurred without gross alteration in glucose homeostasis. These results are in contrast to the results in the 129S2 or F1 hybrid B6129SF1/J groups, both of which exhibited fasting hyperglycemia and improved glucose tolerance with high-dose tamoxifen. In summary, tamoxifen has a cytostatic effect on β-cells in C57Bl/6J mice that appears to be independent of any possible changes in glucose homeostasis.

### Tamoxifen-associated changes in islet gene expression

To further test the impact of tamoxifen we quantified gene expression in islets of treated mice. We isolated islets from F1 hybrid B6129SF1/J mice that were treated with tamoxifen (0.3mg/g of body weight) or oil and then performed RT-PCR using a panel of qPCR primer probe sets focused upon regulators of cell cycle entry (Fig. 6a). Islets from tamoxifen-treated mice exhibited reduced cyclin D1 and D2 RNA expression (cyclin D1 tamoxifen 74% vs. control, p=0.04; cyclin D2 tamoxifen 66% vs. control, p=0.0098)(Fig. 6b, Table S8). In contrast, expression of other cell cycle related genes were not consistently altered by tamoxifen treatment. Cyclins D2 and D1 are known to have essential roles to promote β-cell proliferation in mice [29]. Thus, sharply reduced D-type cyclin RNAs could possibly explain the reduced β-cell proliferation of tamoxifen-treated mice.

**Fig. 6.**
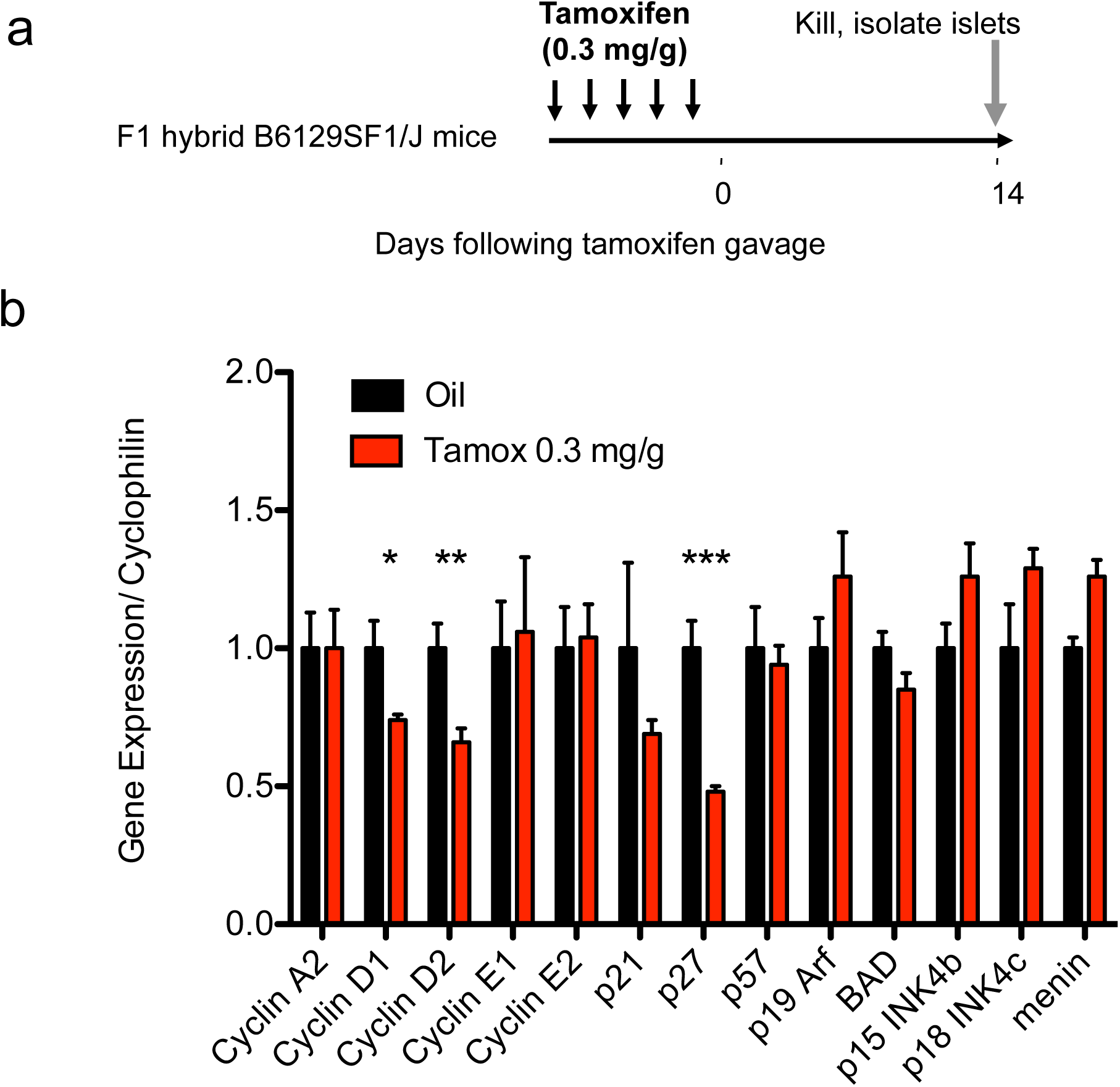
Altered gene expression in islets harvested from tamoxifen-treated mixed genetic background (F1 hybrid B6129SF1/J) mice. (**a**) Schematic indicating time course of treatment. (b) RT-PCR on islet cDNA from tamoxifen-treated mice. Several common genes known to influence cell cycle entry of β-cells were chosen for analysis, including various cyclins, cyclin inhibitors, and tumor suppressors. Results expressed as mean ± SEM for 5 controls and 5 tamoxifen-treated mice per group. *p≤0.05, **p≤0.01, ***p≤0.001, tamoxifen vs. vehicle.

## DISCUSSION

Here we show that tamoxifen inhibits β-cell proliferation in a dose-dependent manner in F1 hybrid B6129SF1/J mice. We further show that high-dose tamoxifen induced fasting hyperglycemia but improved glucose tolerance in mixed genetic background mice. These results are surprising, as tamoxifen has been widely used in the inducible CreERT-loxP system without knowledge of potential toxic effects on β-cell proliferation on non-diabetic mice. These phenomena could complicate β-cell studies that use the inducible Cre^ERT^-loxP system, especially those that employ mice with the C57Bl6/129 or C57Bl/6J genetic backgrounds.

Our study confirms and complements a recently published study by Gannon and colleagues, which similarly communicates unexpected effects of the MIP-CreER transgene and tamoxifen on β-cell growth in C57Bl6/J male mice [42]. Notably, we came across this publication after we had independently planned and completed our work, yet the major conclusions of our studies are similar. Still, our work differs from the Gannon paper in several ways: we carried out dose-response studies using various tamoxifen doses in F1 hybrid B6129SF1/J male mice followed by quantification of β-cell proliferation; we confirmed β-cell proliferation in a cohort with Ki67; we quantified acinar cell turnover; we carried out parallel studies in 129S2 and C57Bl6 to test for genetic background-specific effects of tamoxifen sensitivity; and we performed gene expression studies of islets from tamoxifen treated mice to characterize pathways involved in the β-cell growth. In contrast, Gannon and colleagues also carried out additional studies with high fat diet and with mice containing the MIP-CreER transgene. Taken together, these two independent studies coming to similar conclusions should give investigators great caution before using the inducible Cre^ERT^-loxP system without appropriate controls, including those without tamoxifen.

As expected for a short tern treatment study, β-cell mass was not grossly altered by tamoxifen treatment in F1 hybrid B6129SF1/J mice. BrdU+ β-cells were decreased in high dose tamoxifen treated cohorts: Vehicle treated mice exhibited 12.9% BrdU+ β-cells compared to 4.4% BrdU+ β-cells in mice treated with 0.3mg/g tamoxifen. However, this absolute reduction of 8.5% BrdU+ β-cells would have been expected to result in a net change of only 4% in total β-cells. Thus, the lack of change in β-cell mass was entirely as expected for a short tern study. As a result, we concentrated on quantification of BrdU+ β-cells in our subsequent studies of pure genetic background mice. Similarly, we elected to not quantify β-cell mass in mice that were treated with lower doses of tamoxifen, which had lesser difference in BrdU+ β-cells compared to controls.

The anti-proliferative effects of tamoxifen upon β-cells could be due to estrogen receptor-independent action. In human patients 20-40 mg of tamoxifen is a typical dose for women with breast cancer [19]. 20 mg provides 285 ng/g body weight for a typical 70kg person [19]. Similarly, tamoxifen has potent endogenous estrogen receptor-mediated effects on tissues such as uterus when administered to mice at 100 ng/g body weight [43]. However, typical doses to achieve Cre loxP recombination in transgenic mice are 0.1-0.3 mg/g body weight and therefore 1000-3000x higher than the doses require to activate endogenous estrogen receptors.

Part of the problem with tamoxifen toxicity may be intrinsic to the inducible Cre^ERT^-loxP system. Chambon and colleagues derived the most widely used mutant forms of the estrogen receptor ligand-binding domain and characterized activity of the various mutants when fused to Cre [13]. Their most widely used mutant (Cre-ER^T1^) was insensitive to estradiol but was activated tamoxifen genes [13]. Cre-ER^T1^ requiring 250nM 4-hydroxy-tamoxifen (a high affinity tamoxifen metabolite) to recombine and therefore activate reporter genes [13]. In comparison, native estrogen receptor is activated by sub-nanomolar estradiol and can be efficiently blocked by sub-nanomolar 4-hydroxytamoxifen [44]. The dose of tamoxifen required to efficiently activate Cre^ERT^ and recombine at loxP sites in mice is therefore vastly higher (∼1000 fold) than the amount of tamoxifen necessary to influence endogenous estrogen receptor signals.

Our study has certain limitations. By design we focused on the impact of tamoxifen upon β-cell proliferation in the immediate 2 weeks following drug administration. As a result we cannot determine how long a short course of tamoxifen will impair β-cell proliferation. In addition, we only measured proliferation β-cells and not other islet endocrine cell types. Thus, investigators must consider the possibility that tamoxifen could also negatively impact turnover of other islet endocrine cells of interest.

Tamoxifen-associated changes in β-cell proliferation might be unavoidable. In our studies anti-proliferative effects were present in mice that received tamoxifen at 0.1 to 0.3 mg/g body weight, which correspond to the typical doses used to achieve high-efficiency Cre loxP recombination in transgenic mice. Thus, additional controls are probably necessary for most studies to determine and attempt to limit any potential confounding effects of high dose tamoxifen. Although reducing tamoxifen doses to as low as 0.025 mg/g body weight could be considered to limit the anti-proliferative effects of tamoxifen, this tactic is unlikely to yield sufficient recombination efficiency in β-cells using current technologies [2]. Mice with 129S2 genetic backgrounds might also be deployed to limit the potential effects of tamoxifen on β-cell proliferation, assuming such lines also allow high efficiency Cre-loxP mediated recombination. Alternatively, investigators might simply need to consider and acknowledge the potential anti-proliferative effects of tamoxifen on β-cells within their studies.

The signals that mediate tamoxifen-inhibited β-cell proliferation are unclear. We were able to conduct preliminary experiments with whole islets to interrogate the mechanism of tamoxifen’s toxic effects on β-cell proliferation. These studies indicate that RNAs of both major D-type cyclins in mouse islets (D1 and D2) are altered by tamoxifen administration. These findings are consistent with previous studies implicating cyclins D1 and D2 in β-cell proliferation [29,45,46]. Moreover, the anti-proliferative effects of tamoxifen did not extend to the acinar cell turnover. Still, additional studies are required to determine the role of specific β-cell-mitogenic pathways. Given the effects of high dose tamoxifen upon PKC mitogenic signals [20,21], PKCζ must be considered a top candidate, as PKCζ signals are involved in obesity associated β-cell proliferation via mTOR and Cyclin-D2 [47].

Tamoxifen may have additional effects on mammalian development beyond β-cell proliferation. A study Moon and colleagues involved expression of CreERT via tamoxifen from the Forkhead box protein A2 (Foxa2) locus in mice [48]. Various doses and methods of tamoxifen administration were tested to develop the system. The study noted developmental delay, abnormal head development, and intrauterine hemorrhage in the embryos of pregnant mice that received 0.0625 mg/g intra-peritoneal tamoxifen. In contrast, embryos that received 0.12 mg/g tamoxifen via oral gavage appeared to be normal. Tamoxifen may also influence behavioral phenotyping in tests such as the forced swim test [49]. Of course, as a potent anti-estrogen tamoxifen also has potent effects upon embryonic development and can be associated with fetal loss [50].

In summary, tamoxifen suppressed β-cell proliferation but not acinar proliferation in a dose-dependent manner. Genetic background effects seemed to influence the impact of tamoxifen upon β-cell proliferation, with more potent effects in C57Bl6 mice compared to 129S2 mice. Tamoxifen was associated with modest changes in glucose homeostasis that could be separated from changes in β-cell proliferation. Investigators must judiciously use additional controls to limit potential confounding effects upon β-cell proliferation.

## Acknowledgements

We thank Dr. Peter J. Kushner (Olema Pharmaceuticals Inc., San Francisco, CA) for helpful discussions regarding estrogen receptor biology.

## Data availability

All available data is included in supplemental tables.

## Author Contributions

Conceived and designed the experiments: SHA, AG, MMR, & JAK. Performed the experiments: SHA, AG, MMR, CJL, AJC. Analyzed the data: SHA, AG, MMR, CJL, AJC & JAK. Wrote the paper: SHA & JAK. JAK is the guarantor of this work and, as such, had full access to all the data in the study and takes responsibility for the integrity of the data and the accuracy of the data analysis.

---

**Supplemental Table 1.**
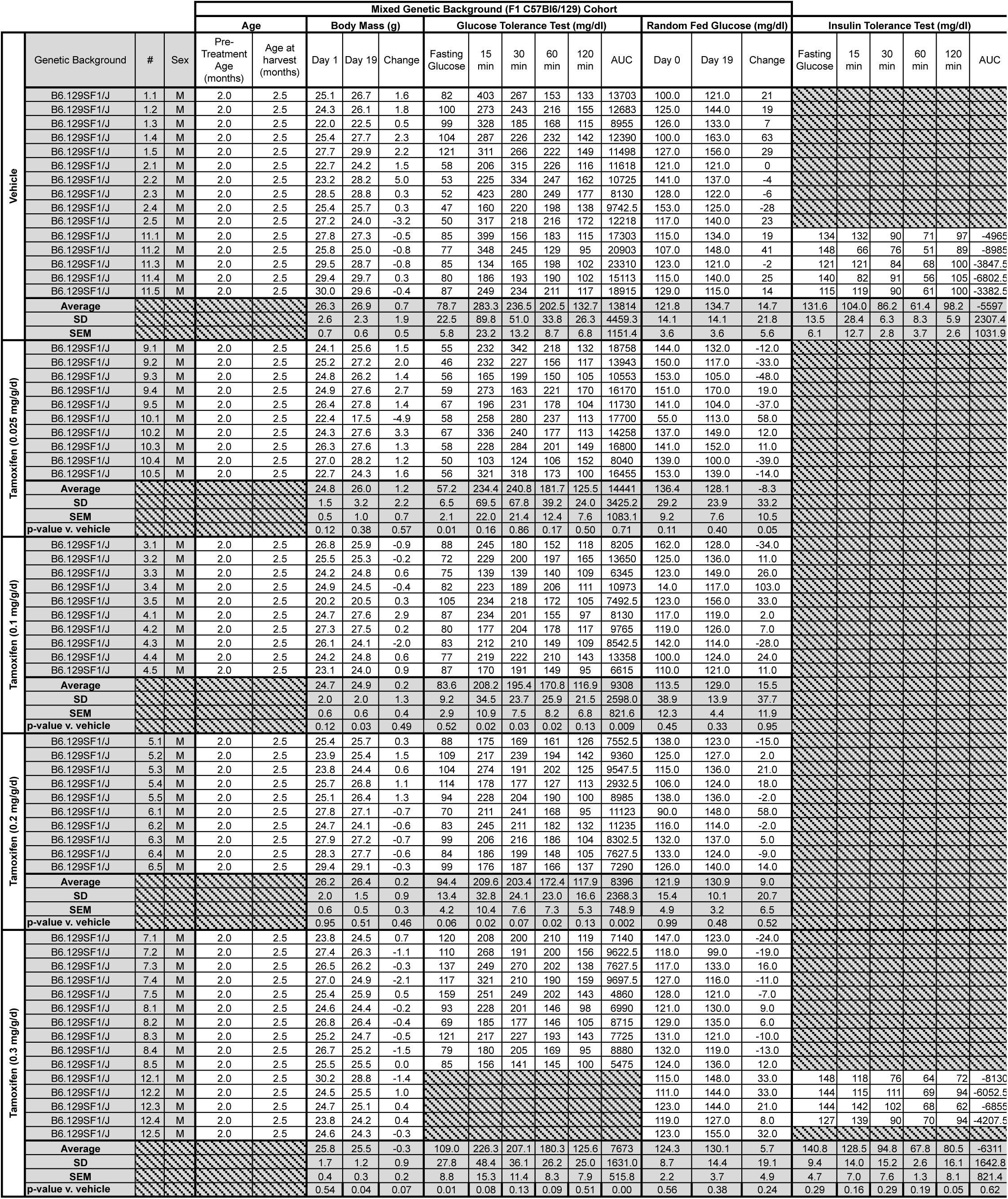
Individual Physiologic Data from Tamoxifen-Treated Mixed Genetic Background (F1 C57Bl6/129) Mice. Genetic Background, Mouse ID, sex, age prior to treatment and at harvest (months), body mass (g) at begining and end of experiment, change in body mass over treatment (g), glucose tolerance test (GTT) blood glucose measurements (mg/dl) after a 16-hour overnight fast, and at time 15, 30, 60, and 120 minutes, area above curve random(mg/dl/min), fed glucose values (mg/dl) prior to treatment (day 0) and at end of treatment (day 19), insulin tolerance test (ITT) 0.8 units per gram of body weight) blood glucose measurements (mg/dl), area above curve random(mg/dl/min).

**Supplemental Table 2.** Individual Beta Cell Morphemtry Data from Tamoxifen-Treated Mixed Genetic Background (F1 C57Bl6/129) Mice. Genetic background, mouse ID, pancreas mass (mg), β-cell area (%), β-cell mass (mg).

| Mixed Genetic Background (F1 C57Bl6/129) Cohort |  |  |  |  |  |
| --- | --- | --- | --- | --- | --- |
| Condition | | | β-cell Morphometry | | |
| | Genetic Background | # | Pancreas Mass (mg) | β-cell area in pancreas (% Total) | β-cell mass (mg) |
| Vehicle | B6.129SF1/J | 2.1 | 280.0 | 0.28 | 0.77 |
|  | B6.129SF1/J | 2.2 | 233.0 | 0.57 | 1.33 |
|  | B6.129SF1/J | 2.3 | 274.0 | 0.59 | 1.62 |
|  | B6.129SF1/J | 2.4 | 295.0 | 0.51 | 1.49 |
|  | B6.129SF1/J | 2.5 | 241.0 | 0.28 | 0.66 |
|  | **Average** |  | 264.6 | 0.44 | 1.17 |
|  | **SD** |  | 26.5 | 0.16 | 0.43 |
|  | **SEM** |  | 11.8 | 0.07 | 0.19 |
| Tamoxifen (0.3 mg/g/d) | B6.129SF1/J | 8.1 | 247.0 | 0.25 | 0.62 |
|  | B6.129SF1/J | 8.2 | 243.0 | 0.40 | 0.98 |
|  | B6.129SF1/J | 8.3 | 175.0 | 0.22 | 0.38 |
|  | B6.129SF1/J | 8.4 | 209.0 | 0.49 | 1.02 |
|  | B6.129SF1/J | 8.5 | 242.0 | 0.48 | 1.17 |
|  | **Average** |  | 223.2 | 0.37 | 0.83 |
|  | **SD** |  | 31.0 | 0.13 | 0.33 |
|  | **SEM** |  | 13.9 | 0.06 | 0.15 |
|  | **p-value v. vehicle** |  | 0.05 | 0.43 | 0.20 |

**Supplemental Table 3.** Individual Beta Cell TUNEL Data from Tamoxifen-Treated Mixed Genetic Background (F1 C57Bl6/129) Mice. Genetic background, mouse ID, β-cells, TUNEL+ DAPI+ Insulin-Islet Cells, TUNEL+ DAPI+ Insulin-Islet Cells (% of β-cells), TUNEL+ β-cells, TUNEL+ β-cells (% of β-cells).

| Mixed Genetic Background (F1 C57Bl6/129) Cohort |  |  |  |  |  |  |  |
| --- | --- | --- | --- | --- | --- | --- | --- |
| Condition | Genetic Background | # | β-cell Morphometry | | | | |
| | | | β-cells | TUNEL+ DAPI+ Insulin- Islet Cells | TUNEL+ DAPI+ Insulin- Islet Cells (% of β-cells) | TUNEL+ β-cells | TUNEL+ β-cells (% of β-cells) |
| Vehicle | B6.129SF1/J | 1.1 | 1147 | 6.0 | 0.00523 | 1.0 | 0.00087 |
|  | B6.129SF1/J | 1.2 | 3252 | 28.0 | 0.00861 | 1.0 | 0.00031 |
|  | B6.129SF1/J | 1.3 | 1629 | 11.0 | 0.00675 | 1.0 | 0.00061 |
|  | B6.129SF1/J | 1.4 | 4955 | 13.0 | 0.00262 | 1.0 | 0.00020 |
|  | B6.129SF1/J | 1.5 | 4388 | 6.0 | 0.00137 | 0.0 | 0.00000 |
|  | B6.129SF1/J | 2.1 | 4253 | 3.0 | 0.00071 | 1.0 | 0.00024 |
|  | B6.129SF1/J | 2.2 | 5883 | 4.0 | 0.00068 | 0.0 | 0.00000 |
|  | B6.129SF1/J | 2.3 | 2499 | 2.0 | 0.00080 | 0.0 | 0.00000 |
|  | B6.129SF1/J | 2.4 | 4475 | 4.0 | 0.00089 | 0.0 | 0.00000 |
|  | B6.129SF1/J | 2.5 | 937 | 10.0 | 0.01067 | 1.0 | 0.00107 |
|  | Average |  | 3342 | 8.7 | 0.00383 | 0.6 | 0.00033 |
|  | SD |  | 1717 | 7.7 | 0.00373 | 0.5 | 0.00039 |
|  | SEM |  | 543 | 2.4 | 0.00118 | 0.2 | 0.00012 |
| Tamoxifen (0.3 mg/g/d) | B6.129SF1/J | 7.1 | 4382 | 4.0 | 0.00091 | 0.0 | 0.00000 |
|  | B6.129SF1/J | 7.2 | 1943 | 48.0 | 0.02470 | 1.0 | 0.00051 |
|  | B6.129SF1/J | 7.3 | 5988 | 28.0 | 0.00468 | 2.0 | 0.00033 |
|  | B6.129SF1/J | 7.4 | 5686 | 8.0 | 0.00141 | 0.0 | 0.00000 |
|  | B6.129SF1/J | 7.5 | 1061 | 27.0 | 0.02545 | 1.0 | 0.00094 |
|  | B6.129SF1/J | 8.1 | 924 | 7.0 | 0.00758 | 0.0 | 0.00000 |
|  | B6.129SF1/J | 8.2 | 791 | 7.0 | 0.00885 | 0.0 | 0.00000 |
|  | B6.129SF1/J | 8.3 | 3928 | 39.0 | 0.00993 | 1.0 | 0.00025 |
|  | B6.129SF1/J | 8.4 | 3133 | 22.0 | 0.00702 | 0.0 | 0.00000 |
|  | B6.129SF1/J | 8.5 | 4083 | 8.0 | 0.00196 | 0.0 | 0.00000 |
|  | Average |  | 3192 | 19.8 | 0.00925 | 0.5 | 0.00020 |
|  | SD |  | 1938 | 15.4 | 0.00891 | 0.7 | 0.00032 |
|  | SEM |  | 613 | 4.9 | 0.00282 | 0.2 | 0.00010 |
|  | p-value v. vehicle |  | 0.86 | 0.06 | 0.09323 | 0.72 | 0.44289 |

**Supplemental Table 4.**
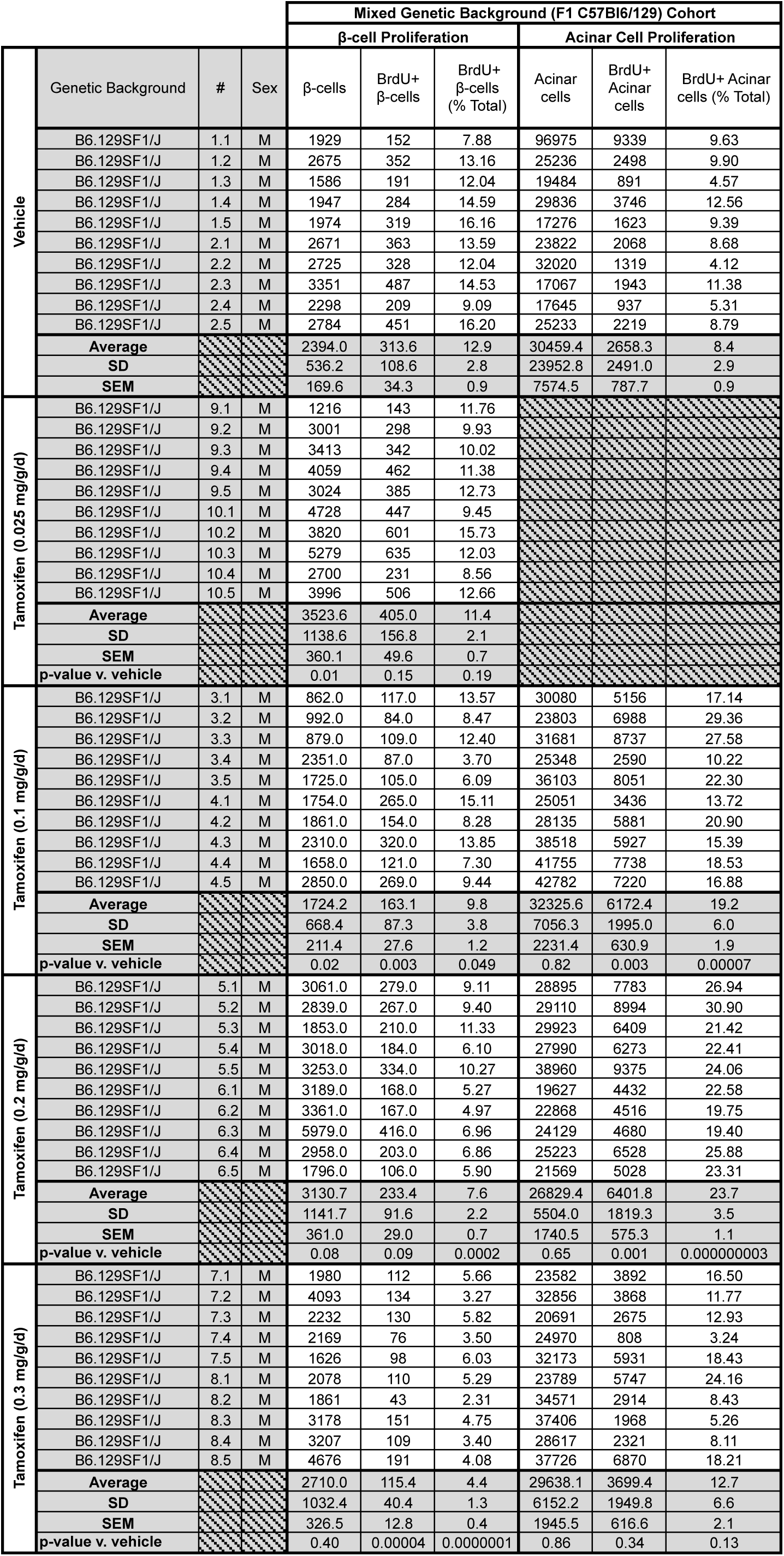
Individual Beta Cell and Acinar Cell Proliferation from Tamoxifen-Treated Mixed Genetic Background (F1 C57Bl6/129) Mice. Mouse ID, β-cell (number), BrdU+ β-cells (number; % total), acinar cell (number), BrdU+ acinar cells (number; % total).

**Supplemental Table 5.** Individual Beta Cell Ki67 Proliferation from Tamoxifen-Treated Mixed Genetic Background (F1 C57Bl6/129) Mice. Mouse ID, β-cell (number), Ki67+ β-cells (number; % total).

| Mixed Genetic Background (F1 C57Bl6/129) Cohort |  |  |  |  |  |  |
| --- | --- | --- | --- | --- | --- | --- |
| Condition | | | | β-cell Proliferation | | |
| | Genetic Background | # | Sex | β-cells | Ki67+ β-cells | Ki67+ β-cells (% Total) |
| Vehicle | B6.129SF1/J | 1.1 | M | 2450 | 20 | 0.82 |
|  | B6.129SF1/J | 1.2 | M | 4991 | 50 | 1.00 |
|  | B6.129SF1/J | 1.3 | M | 1946 | 11 | 0.57 |
|  | B6.129SF1/J | 1.4 | M | 2802 | 48 | 1.71 |
|  | B6.129SF1/J | 2.1 | M | 2450 | 20 | 0.82 |
|  | B6.129SF1/J | 2.2 | M | 4991 | 50 | 1.00 |
|  | B6.129SF1/J | 2.3 | M | 1946 | 11 | 0.57 |
|  | Average |  |  | 3082.3 | 30.0 | 0.93 |
|  | SD |  |  | 1338.3 | 18.5 | 0.39 |
|  | SEM |  |  | 505.8 | 7.0 | 0.15 |
| Tamoxifen (0.3 mg/g/d) | B6.129SF1/J | 7.1 | M | 1986 | 11 | 0.25 |
|  | B6.129SF1/J | 7.2 | M | 1548 | 1 | 0.40 |
|  | B6.129SF1/J | 7.3 | M | 3002 | 19 | 0.22 |
|  | B6.129SF1/J | 8.2 | M | 1796 | 5 | 0.40 |
|  | B6.129SF1/J | 8.4 | M | 3268 | 14 | 0.49 |
|  | B6.129SF1/J | 8.5 | M | 4424 | 23 | 0.48 |
|  | Average |  |  | 2670.7 | 12.2 | 0.37 |
|  | SD |  |  | 1098.6 | 8.3 | 0.11 |
|  | SEM |  |  | 448.5 | 3.4 | 0.05 |
|  | p-value v. vehicle |  |  | 0.56 | 0.052 | 0.007 |

**Supplemental Table 6.**
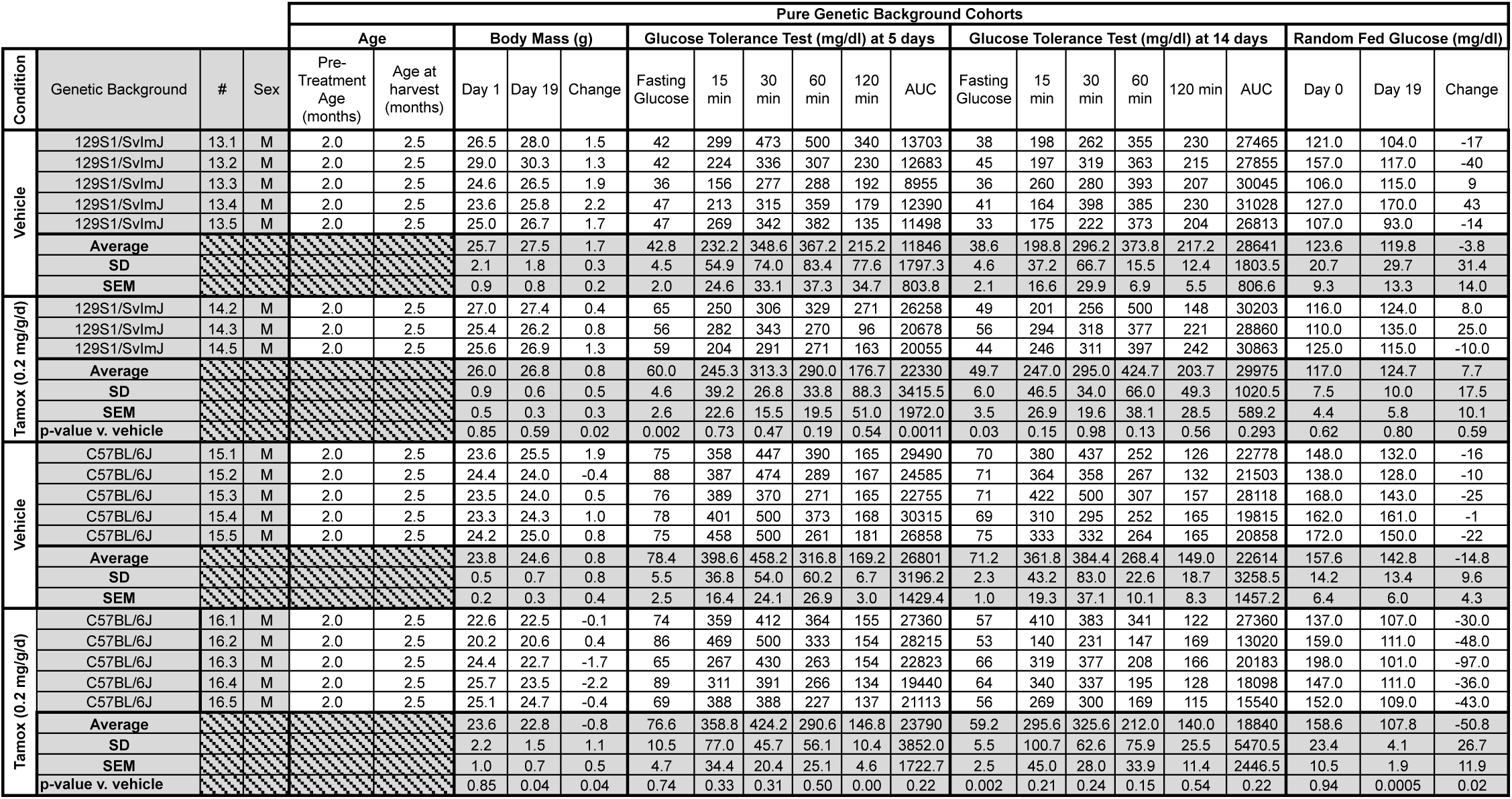
Individual Physiologic Data from Tamoxifen-Treated Pure Genetic Background (129S1/SvImJ or C57BL/6J) Mice. Genetic Background, Mouse ID, sex, age prior to treatment and at harvest (months), body mass (g) at begining and end of experiment, change in body mass over treatment (g), glucose tolerance tests blood glucose measurements (mg/dl) after a 16-hour overnight fast, and at time 15, 30, 60, and 120 minutes, random fed glucose values (mg/dl) prior to treatment (day 0) and at end of treatment (day 19).

**Supplemental Table 7.** Individual Beta Cell Proliferation from Tamoxifen-Treated Pure Genetic Background (129S1/SvImJ or C57BL/6J) Mice. Mouse ID, β-cell (number), BrdU+ β-cells (number; % total).

| Pure Genetic Background Cohorts |  |  |  |  |  |  |
| --- | --- | --- | --- | --- | --- | --- |
| Condition | | | | β-cell Proliferation | | |
| | Genetic Background | # | Sex | β-cells | BrdU+ β-cells | BrdU+ β-cells (% Total) |
| Vehicle | 129S1/SvImJ | 13.1 | M | 1981 | 244 | 12.32 |
|  | 129S1/SvImJ | 13.2 | M | 870 | 73 | 8.39 |
|  | 129S1/SvImJ | 13.3 | M | 1804 | 243 | 13.47 |
|  | 129S1/SvImJ | 13.4 | M | 1048 | 111 | 10.59 |
|  | 129S1/SvImJ | 13.5 | M | 567 | 61 | 10.76 |
|  | **Average** |  |  | 1254.0 | 146.4 | 11.11 |
|  | **SD** |  |  | 610.9 | 90.5 | 1.92 |
|  | **SEM** |  |  | 273.2 | 40.5 | 0.86 |
| Tamox (0.2 mg/g/d) | 129S1/SvImJ | 14.2 | M | 807 | 60 | 7.43 |
|  | 129S1/SvImJ | 14.3 | M | 3161 | 349 | 11.04 |
|  | 129S1/SvImJ | 14.5 | M | 1276 | 115 | 9.01 |
|  | **Average** |  |  | 1748.0 | 174.7 | 9.16 |
|  | **SD** |  |  | 1246.0 | 153.5 | 1.81 |
|  | **SEM** |  |  | 719.4 | 88.6 | 1.04 |
|  | **p-value v. vehicle** |  |  | 0.47 | 0.75 | 0.21 |
| Vehicle | C57BL/6J | 15.1 | M | 3304 | 303 | 9.17 |
|  | C57BL/6J | 15.2 | M | 1493 | 121 | 8.10 |
|  | C57BL/6J | 15.3 | M | 1043 | 51 | 4.89 |
|  | C57BL/6J | 15.4 | M | 966 | 66 | 6.83 |
|  | C57BL/6J | 15.5 | M | 2375 | 155 | 6.53 |
|  | **Average** |  |  | 1836.2 | 139.2 | 7.10 |
|  | **SD** |  |  | 993.6 | 100.7 | 1.63 |
|  | **SEM** |  |  | 444.4 | 45.0 | 0.73 |
| Tamox (0.2 mg/g/d) | C57BL/6J | 16.1 | M | 3601 | 89 | 2.47 |
|  | C57BL/6J | 16.2 | M | 2452 | 81 | 3.30 |
|  | C57BL/6J | 16.3 | M | 1784 | 22 | 1.23 |
|  | C57BL/6J | 16.4 | M | 414 | 14 | 3.38 |
|  | C57BL/6J | 16.5 | M | 233 | 10 | 4.29 |
|  | **Average** |  |  | 1696.8 | 43.2 | 2.9 |
|  | **SD** |  |  | 1413.5 | 38.5 | 1.1 |
|  | **SEM** |  |  | 632.1 | 17.2 | 0.5 |
|  | **p-value v. vehicle** |  |  | 0.86 | 0.08 | 0.00158 |

**Supplemental Table 8.**
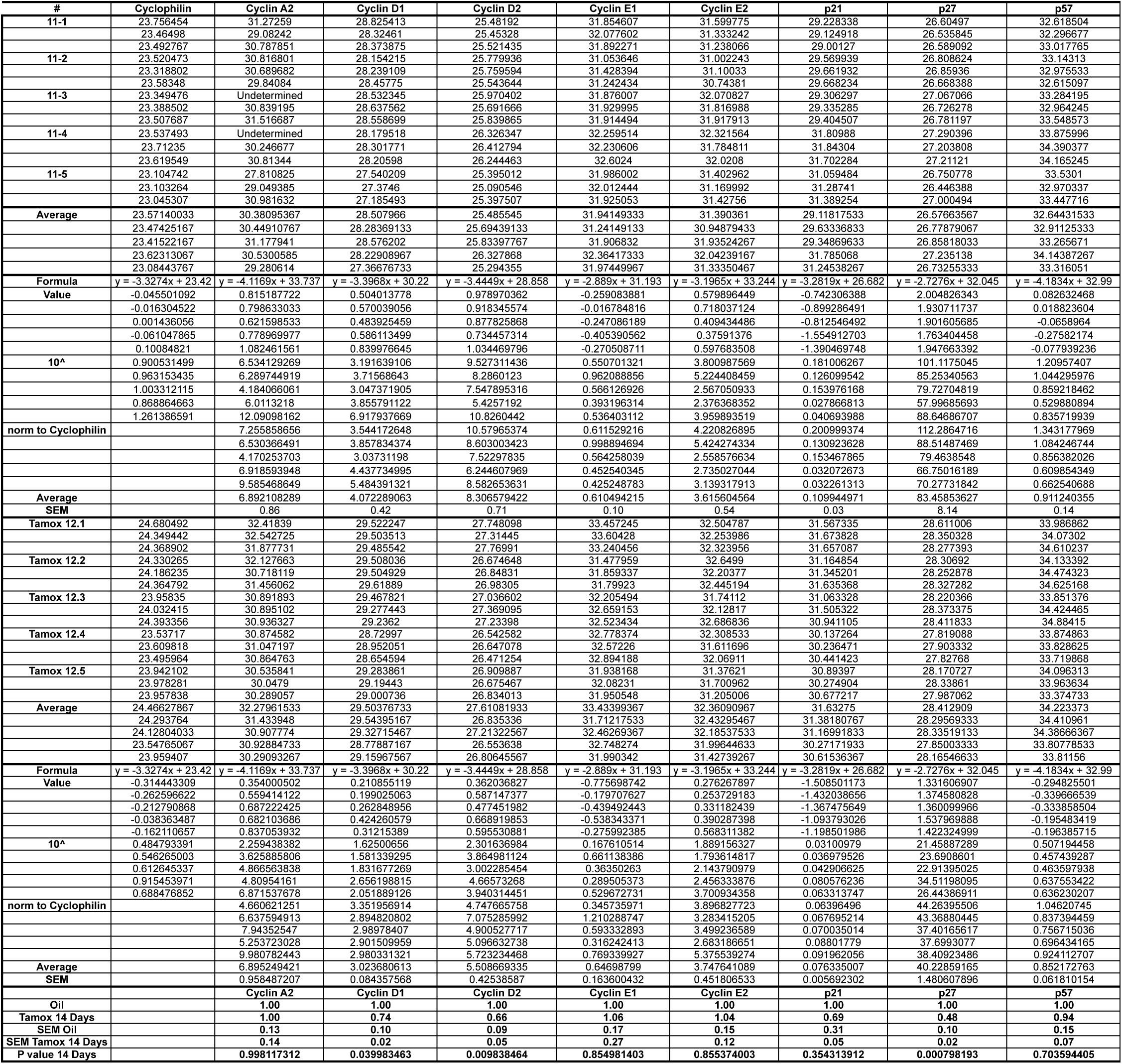
Quantitative RT-PCR: Gene Expression Studies of Islets Harvested From Tamoxifen Treated Mixed Genetic Background (F1 C57Bl6/129) Mice. Mouse ID, cDNAs detected (cyclophillin, cyclin A2, cyclin D1,cyclin D2, cyclin D3, cyclin E1, cylin E2, p21, p27, p57, p19 Arf, p15 INK4b, p18 INK4c, menin)

